# Predictors of virus prevalence and diversity across a wild bumblebee community

**DOI:** 10.1101/2021.01.06.425554

**Authors:** David J. Pascall, Matthew C. Tinsley, Bethany L. Clark, Darren J. Obbard, Lena Wilfert

## Abstract

Viruses are key regulators of natural populations. Despite this, we have limited knowledge of the diversity and ecology of viruses that lack obvious fitness effects on their host. This is even the case in wild host populations that provide ecosystem services, where small fitness effects may have major ecological and financial impacts in aggregate. One such group of hosts are the bumblebees, which have a major role in the pollination of food crops and have suffered population declines and range contractions in recent decades. In this study, we used a multivariate generalised linear mixed model to investigate the ecological factors that determine the prevalence of four recently discovered bumblebee viruses (Mayfield virus 1, Mayfield virus 2, River Liunaeg virus and Loch Morlich virus), and two previously known viruses that infect both wild bumblebees and managed honeybees (Acute bee paralysis virus and Slow bee paralysis virus). We show that the recently discovered bumblebee viruses were more genetically diverse than the viruses shared with honeybees, potentially due to spillover dynamics of shared viruses. We found evidence for ecological drivers of prevalence in our samples, with relatively weak evidence for a positive effect of precipitation on the prevalence of River Luinaeg virus. Coinfection is potentially important in shaping prevalence: we found a strong positive association between River Liunaeg virus and Loch Morlich virus presence after controlling for host species, location and other relevant ecological variables. This study represents a first step in the description of predictors of bumblebee infection in the wild not driven by spillover from honeybees.

## Introduction

Viruses are among the most abundant and diverse groups of organisms on Earth (Wommack et al., 2015); wherever they are looked for, they are found in other species as obligate pathogens. Despite this, viral ecology in natural populations remains understudied. In the wild, infection is generally only recorded when clear symptoms of the underlying disease are present, such as discolouration, aberrant tissue structures, or an increase in mortality. However, these symptoms are rarely detected in natural infections (Mackenzie & Jeggo, 2013). By focusing only on those viruses that cause obvious symptoms in well-studied host species, we are likely to be underestimating the diversity of viruses and their ecological importance in regulating natural populations. For example, the dynamics of algal blooms are strongly driven by density dependent regulation of the algae through viral infections (Bratbak et al., 1991; Brussaard et al., 1996). The development of relatively cheap and easily applied molecular techniques has allowed the detection and identification of potentially pathogenic organisms within both the host and the environment, enabling the systematic study of viral ecology in wild populations (e.g. (Webster et al., 2015)). This is especially important for threatened host species, where understanding the viral burden may have conservation implications (Gordon et al., 2015).

Pollinators are economically important and so their viruses are important in turn. Over 50 viruses have now been described in bees, and their importance to survival is well recognised (e.g. (McMahon et al., 2018)). However, the majority of this work has been performed in the European honeybee, *Apis mellifera*, thus the knowledge of the viral ecology of bumblebees is more limited. Some honeybee work is transferable to bumblebees; for instance, viruses known from honeybees have pathogenic effects in the buff-tailed bumblebee, *Bombus terrestris* (Fürst et al., 2014; Graystock et al., 2016; Manley et al., 2017), and their prevalences have been assayed across the UK (Fürst et al., 2014; McMahon et al., 2015). However, few predictors of infection in bumblebees other than the presence of sympatric honeybees have been described in any depth.

In the wild, differences in viral prevalence between hosts or locations can be explained by a variety of ecological factors. If a virus is spread by environmental contamination or aerosolisation, then abiotic factors can be important. In bumblebees, infection is often thought to take place at flowers (Durrer & Schmid-Hempel, 1994; Graystock et al., 2015; McArt et al., 2014) and so factors that reduce contamination of floral structures may be predicted to reduce the rate of infection in the general bumblebee population (Adler et al., 2020); obvious mechanisms are viral deactivation, flower visitation rates and physical cleaning. The rate of viral deactivation can be increased in high temperatures, both independently and through an interaction with relative humidity (Mbithi et al., 1991). Additionally, high UV levels may deactivate virus particles rapidly (Lytle & Sagripanti, 2005), an effect thought to be highly important in the regulation of viral populations in oceanic waters (Suttle & Chen, 1992).Furthermore, bees must physically reach the flowers where infection can occur, so factors that change the rate of contact of workers with heavily contaminated flowers may also modify viral prevalence. Wind speed affects the relative rates of pollen and nectar collection (Peat & Goulson, 2005), which may alter flower visitation and the energetic costs of foraging (T. J. Wolf et al., 1999), consequently affecting susceptibility to infection. Precipitation can also limit bumblebee foraging and therefore flower contamination risk (Peat & Goulson, 2005). Finally, heavy rain and strong winds may physically clean the flowers. However, environmental conditions would only be expected to lead to interspecific prevalence differences locally through species-specific effects on bee behaviour.

Another factor that can explain infection in host populations is the presence of other infectious agents. Co-occurrences of particular pathogens have been observed in many species and can drive infection dynamics with certain combinations being over- or under-represented relative to chance expectations (Johnson et al., 2015; Tollenaere et al., 2016). This may be due to synergistic or antagonistic interactions between the pathogens. For example, mechanical damage to tissues by a primary infection allowing easier access for secondary pathogens (Joseph et al., 2013) or due to competitive exclusion of pathogens with the same niche within the host (Amaku et al., 2013). Very few studies in pollinators have looked for these between pathogen interactions in a statistically rigorous manner.

Given the complexity of the pollinator system, it is also unclear how much genetic diversity will be present in the viral populations. As some viruses such as the re-emerging Deformed wing virus represent spillovers from honeybee populations into wild pollinator populations (Fürst et al., 2014; Manley et al., 2019; Wilfert et al., 2016), the genetic diversity in wild pollinators for these viruses will likely represent a potentially non-random sample of viral diversity in its maintenance host. For viruses that are maintained in bumblebees, however, the diversity would be expected to be impacted by the normal evolutionary patterns of mutation, selection and drift. Depending on the population size and the historical selection regime, this could lead to very high diversity, or almost none if a major selective event or bottleneck had recently occurred.

Here we present an exploratory investigation into the determinants of viral prevalence and genetic diversity in wild bumblebee populations consisting of 13 species from nine sites across Scotland. We explore the effect of differences in temperature, UV radiation, wind speed and precipitation on the prevalences of four recently discovered bumblebee viruses Mayfield virus 1 (MV1), Mayfield virus 2 (MV2), River Liunaeg virus (RLV), and Loch Morlich virus (LMV), where fitness effects have not yet been tested. We also explore whether or not there are observable statistical interactions between the pathogens that might indicate facilitative or suppressive interactions. We consider the genetic diversity in these viruses and contrast this with two viruses known from honeybees, Acute bee paralysis virus (ABPV) and Slow bee paralysis virus (SBPV). We show that the viruses described only in bumblebees are universally more diverse than ABPV and SBPV and that there is a strong positive association between LMV and RLV infection.

## Methods

### Sampling Regime and Molecular Work

Samples were derived from the field collections described in Pascall et al. (2018). Briefly, we collected a total of 759 bumblebees of 13 species from 9 sites across Scotland, UK (Supp Table 1; Fig. 1). The Ochil Hills, Glenmore, Dalwhinnie, Stirling, Iona, Staffa and the Pentlands were sampled in 2009, while Edinburgh and Gorebridge were sampled in 2011. We performed individual RNA extractions using TRIzol (Life Technologies) following the manufacturers’ standard protocol. RNA was transcribed into cDNA using random hexamers and goScript MMLV reverse transcriptase (Promega) following the manufacturers’ instructions.

**Table 1.** The Morisita-Horn dissimilarities of the bumblebee compositions of the different sampling sites. 90% shortest posterior density intervals for the index are in brackets.

|  | Dalwhinnie | Edinburgh | Glenmore | Gorebridge | Iona | Ochils | Pentlands | Staffa | Stirling |
| --- | --- | --- | --- | --- | --- | --- | --- | --- | --- |
| Dalwhinnie | 0 |  |  |  |  |  |  |  |  |
| Edinburgh | 0.764<br>(0.548,<br>0.909) | 0 |  |  |  |  |  |  |  |
| Glenmore | 0.491<br>(0.228,<br>0.774) | 0.934<br>(0.855,<br>0.971) | 0 |  |  |  |  |  |  |
| Gorebridge | 0.548<br>(0.355,<br>0.733) | 0.162<br>(0.104,<br>0.217) | 0.918<br>(0.840,<br>0.964) | 0 |  |  |  |  |  |
| Iona | 0.220<br>(0.079,<br>0.501) | 0.653<br>(0.494,<br>0.819) | 0.817<br>(0.629,<br>0.914) | 0.415<br>(0.278,<br>0.532) | 0 |  |  |  |  |
| Ochils | 0.741<br>(0.527,<br>0.894) | 0.233<br>(0.150,<br>0.325) | 0.920<br>(0.837,<br>0.962) | 0.315<br>(0.243,<br>0.415) | 0.569<br>(0.336,<br>0.737) | 0 |  |  |  |
| Pentlands | 0.591<br>(0.357,<br>0.795) | 0.501<br>(0.278,<br>0.674) | 0.849<br>(0.661,<br>0.934) | 0.440<br>(0.272,<br>0.663) | 0.517<br>(0.324,<br>0.748) | 0.394<br>(0.160,<br>0.572) | 0 |  |  |
| Staffa | 0.491<br>(0.190,<br>0.742) | 0.893<br>(0.780,<br>0.952) | 0.025<br>(0.004,<br>0.104) | 0.898<br>(0.802,<br>0.959) | 0.831<br>(0.602,<br>0.912) | 0.881<br>(0.754,<br>0.943) | 0.782<br>(0.572,<br>0.898) | 0 |  |
| Stirling | 0.780<br>(0.502,<br>0.879) | 0.070<br>(0.019,<br>0.144) | 0.907<br>(0.806,<br>0.964) | 0.174<br>(0.092,<br>0.298) | 0.637<br>(0.451,<br>0.812) | 0.213<br>(0.105,<br>0.357) | 0.450<br>(0.242,<br>0.676) | 0.853<br>(0.721,<br>0.939) | 0 |

**Figure 1.**
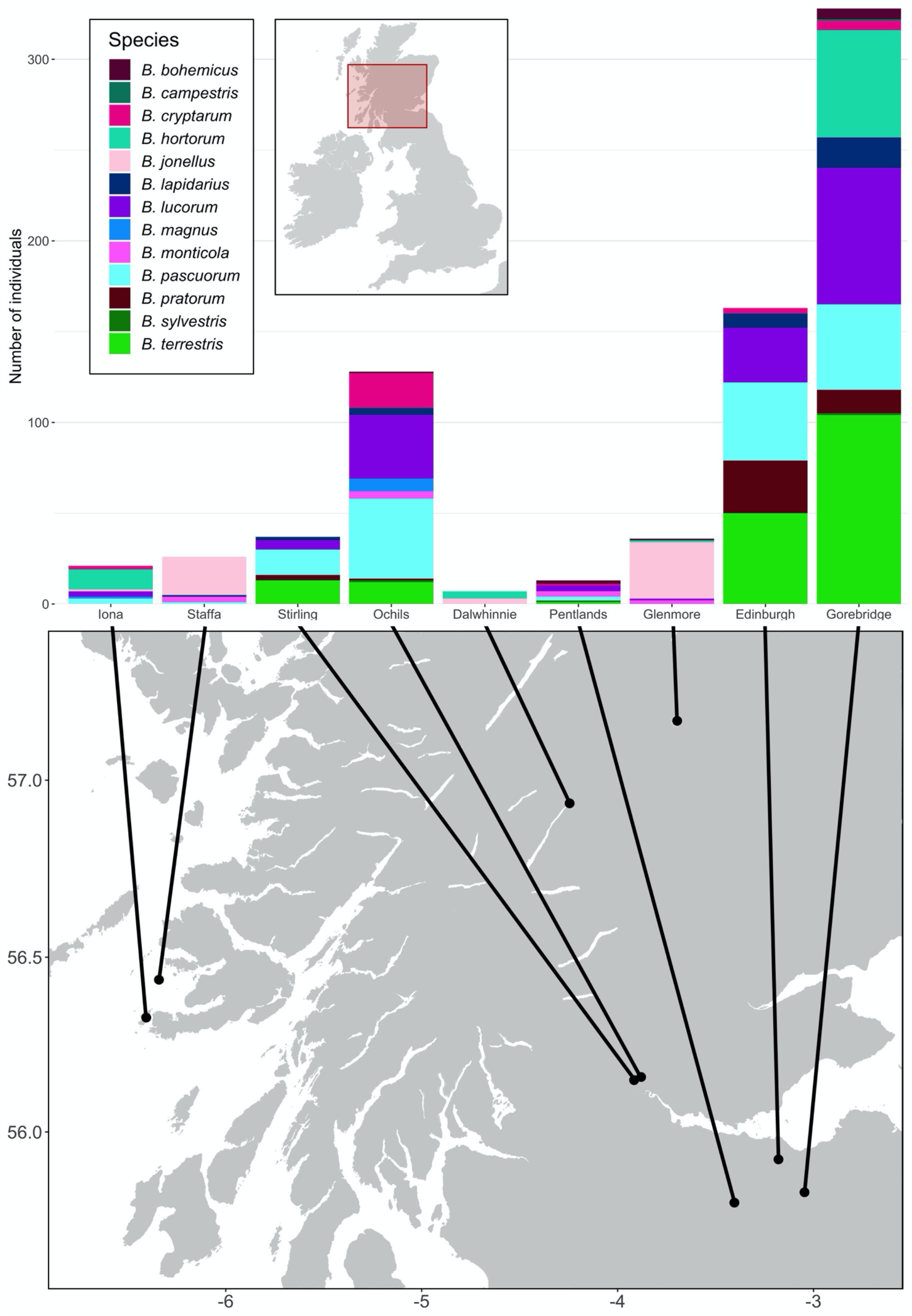
The locations of the sampling sites and species distributions and sample sizes of the bumblebees caught at them. Map adapted from tiles by Stamen Design, under Creative Commons (CC BY 3.0) using data by OpenStreetMap, under the Open Database License.

In this study, we assayed the prevalence of Mayfield virus 1 (MV1), Mayfield virus 2 (MV2), River Luinaeg virus (RLV), Loch Morlich virus (LMV) at the individual level by RTPCR (Supp Table 2). We tested a subset of the samples for Slow bee paralysis virus (SBPV) (n = 544) and Acute bee paralysis virus (ABPV) (n = 385).

**Table 2.** The posterior means and 90% shortest posterior intervals of the coefficients of the effect of each environmental covariate on each virus on the link scale from the multivariate probit model.

|  | Precipitation | Maximum<br>Temperature | Wind Speed |
| --- | --- | --- | --- |
| **River Luinaeg**<br>**virus** | 0.60<br>(0.00, 1.19) | -0.36<br>(-0.99, 0.23) | -0.22<br>(-0.79, 0.33) |
| **Loch Morlich**<br>**virus** | 0.37<br>(-0.46, 1.18) | -0.18<br>(-1.01, 0.70) | -0.02<br>(-0.87, 0.83) |
| **Mayfield virus 1** | -0.50<br>(-1.11, 0.11) | 0.66<br>(-0.11, 1.39) | 0.56<br>(-0.08, 1.17) |
| **Mayfield virus 2** | -0.13<br>(-0.90, 0.63) | -0.10<br>(-0.86, 0.69) | -0.20<br>(-0.95, 0.53) |
| **SBPV** | -0.03<br>(-0.70, 0.66) | 0.11<br>(-0.64, 0.88) | -0.64<br>(-1.40, 0.08) |
| **ABPV** | 0.01<br>(-0.88, 0.91) | -0.15<br>(-1.13, 0.80) | -0.04<br>(-1.00, 0.91) |

### Community Similarity

To estimate host community similarity between sampling sites, we estimated the underlying sampling probability of each bumblebee species at each site by treating the observed samples as being drawn from a multinomial distribution with 24 categories, corresponding to the 24 bumblebee species in the United Kingdom. We use a Dirichlet prior with these 24 categories and a concentration parameter of 1 for each category, implying complete uncertainty about the underlying probability. This has the advantage that the posterior has a known analytical form. Probabilities were estimated independently for each site. Ten thousand simulations were taken from the posterior distributions generated for each site to generate possible values of the underlying sampling probabilities of each bumblebee species at each site, which we assume to be roughly equivalent to the frequency of that bumblebee species at that site. For each of the 10,000 simulations from the posteriors at the sites, we generated estimates of the community dissimilarity using the Morisita-Horn index (Horn, 1966), implemented in the R package vegan (Oksanen et al., 2017). The posterior mode and 90% shortest probability intervals for the dissimilarity index were then reported.

### Prevalence and Climatic Association

Climatic data for each of the nine sites at which bees were collected was taken from the WorldClim database at 1km resolution (Fick & Hijmans, 2017). Predictions for July and August derived from data from 1960-2010 were extracted for mean daily maximum temperature, mean precipitation, mean solar radiation and mean wind speed at the grid reference for the sites with a buffer area of 2km to account for the fact that bumblebees forage over approximately that distance (Wolf & Moritz, 2008). All values were averaged to generate a consensus value for that site and then mean-centred and scaled to unit standard deviation. At this point, we tested the correlation between the variables. Mean solar radiation and mean daily maximum temperature were highly correlated (Pearson correlation: 0.78), so only mean maximum temperature was carried forward. All remaining variables had low Pearson correlations between them in the range of −0.4 to 0.4.

We tested associations between individual prevalence and climate data using Stan version 2.18.2 (Carpenter et al., 2017) via the rstan interface (Stan Development Team, 2016) in R version 3.6 (R Core Development Team, 2016). A multivariate probit model was fitted, with random host, location and host-location effects, and mean maximum temperature, precipitation, and wind speed as fixed effects for each virus. The usage of the multivariate probit allows us to test for excess co-infection between viruses in the study, i.e. coinfection beyond random expectations after the effect of shared covariates has been removed. As the number of sampling locations was small, we expected our ability to accurately determine the size and direction of effects caused by ecological covariates would be limited. To reduce the effect of drawing spurious conclusions due to our small number of sites, we applied regularisation as recommended by Lemoine et al. (2016), using a regularising prior distribution. The global intercept for each virus was given a Gaussian (mu=0, sigma=10) prior, which does not substantially penalise low probabilities. Each fixed effect coefficient was given a Gaussian (mu=0, sigma=1) prior, which, given that the fixed effects act at the site level, should dominate the likelihood if the effect is small. Random effects were drawn from normal distributions centred at 0 with estimated standard deviations. In all cases, the standard deviations were given Exponential (lambda=2) hyperpriors, which are only weakly informative on the logit scale when the data is informative for the standard deviation. The correlation in residuals for the multivariate normal was given a near flat prior using a Lewandoski, Kurowicka and Joe (eta=1) prior. While the Stan code used can regularly give outputs with divergent transitions, the presented model had no divergent transitions over 24000 samples, tail and bulk effective sample sizes of over 400 for all parameters and no Bayesian fraction of missing information warning. We did not perform model selection given our regularising priors, and statements are made based on estimates from the full model.

### Diversity Analysis

To analyse sequence diversity, we used the raw reads from the RNA sequencing described in Pascall et al. (2018). Briefly, these consist of 100bp-paired end RNA-Seq data from pools of *B. terrestris, Bombus pascuorum* and *Bombus lucorum*, each sequenced twice, once using duplex specific normalisation and once using poly-A selection, and a pool of mixed *Bombus* species, sequenced only with poly-A selection. MV1, MV2, RLV, LMV, SBPV Rothamsted (EU035616.1) and ABPV (AF486072.2) sequences were aligned on the TranslatorX server (Abascal et al., 2010), using its MAFFT setting (Katoh & Standley, 2013). Post-alignment, we manually trimmed sequences to the conserved region of the RdRp gene, minus eight codons, owing to the shortness of the RLV sequence. Trailing regions of 200 base pairs at both ends were retained so that reads were not prevented from mapping due to an overhang. This gave final sequence lengths of 1483, 1483, 1536, 1501, 1519 and 1522 base pairs for MV1, MV2, RLV, LMV, SBPV and ABPV respectively. Raw bioinformatic reads were trimmed in sickle version 1.33 using the default parameters (Joshi & Fass, 2011). Overlapping mate reads were combined by FLASH version 1.2.11 using the default settings (Magoč & Salzberg, 2011). Reads were aligned to the RdRp sequences generated above using MOSAIK version 2.1.73 (Lee et al., 2014). Both merged reads and singletons from the sickle run were aligned together in the single end setting. Unmerged paired end reads were separately aligned using the paired end setting. In both cases, a quality threshold of 30 was used to remove ambiguously mapping reads. SAM files were recombined after the fact using SAMtools version 1.5 (Li et al., 2009). Given the high coverage of SBPV, MV1 and MV2 duplicate sequences were not marked, as per best practice (McKenna et al., 2010). Variants were called using the default settings in LoFreq* version 1.2.1 (Wilm et al., 2012). Base quality scores were recalibrated using the outputted vcf file in GATK (DePristo et al., 2011). Variant calling and recalibration were repeatedly performed until the base quality scores converged to a stable distribution (a total of four recalibrations). Once the score distribution stabilised, variant calling was performed to generate a set of variants for the entire sample. These variants were used to recalibrate the scores of each species-specific mapping and generate species level variant calls. If the median depth over called differences from the consensus was less than 20, species-virus combinations were removed from the variant analysis. *B. lucorum* was analysed for SBPV, ABPV and MV1. *B. terrestris* was analysed for SBPV, MV1 and MV2. *B. pascuorum* was analysed for ABPV, SBPV and MV2. The mixed *Bombus* pool was analysed for all six viruses.

The number of polymorphic sites was calculated for each virus. Variants with allele frequencies of 1 were removed as these represent fixed differences from the underlying reference sequence. To measure genetic diversity, we used these counts to approximate Watterson’s theta (Watterson, 1975) for each host-virus combination. We had to account for the fact that the make-up of the pools was not precisely the same as the samples that were tested by PCR. As such, we predicted the status of each of the untested individuals in the pools from the model discussed above and took the median and a 90% central credible interval of the number of additional positives (over those confirmed by PCR). We then used these numbers to give bounds on the approximation to Watterson’s theta estimator, by looking at the value of the estimator at those three estimates of the number of infected individuals. The equation used was:

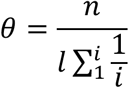

where *θ* is the approximation to Watterson’s theta, *n* is the number of variants, *l* is the length of the sequence and *i* is the number of PCR positives. This method makes two strong assumptions: 1) there is no mixed infection of viral variants in individuals (i.e. that one extra individual represents a single extra count for the harmonic partial sum in the denominator of Watterson’s theta) and 2) all variants present are detectable. The impact of deviation from the first assumption is likely to be small. The marginal change in the partial sum in the denominator decreases with every extra count, so a few missed counts will result in little change to the resultant estimate. The second assumption is more influential, given the larger impact that a missing variant has on the generated number. Given this, we acknowledge that our presented estimates may be conservative.

## Results

### Bumblebee Community Similarity

There were obvious differences in bumblebee community structure between our Scottish sampling sites. The locations we sampled in the south had *B. terrestris, B. pascuorum* and *B. lucorum* dominated communities, whereas those further north had *Bombus jonellus* and *Bombus hortorum* dominated communities (Figure 1, Table 2). Even with its small sample size of 13 bees, the Pentlands, a range of hills in southern Scotland (Figure 1), appeared to represent a third type of community: the presence of *Bombus monticola*, otherwise only found in the highland sites, and an equivalent frequency of *B. pascuorum* and *B. lucorum* makes the community look like a blend of the other community types. This is potentially due to its higher elevation and habitat being similar to the north with large numbers of heathland plants while being situated in the south.

### Prevalence

There were large differences in prevalence of the viruses, all of which are +ssRNA picorna-like viruses, between sites (Figure 2). When broken down to the specific host-location level, sample sizes for many species become small, so the uncertainty around the modal prevalences is correspondingly large.

**Figure 2.**
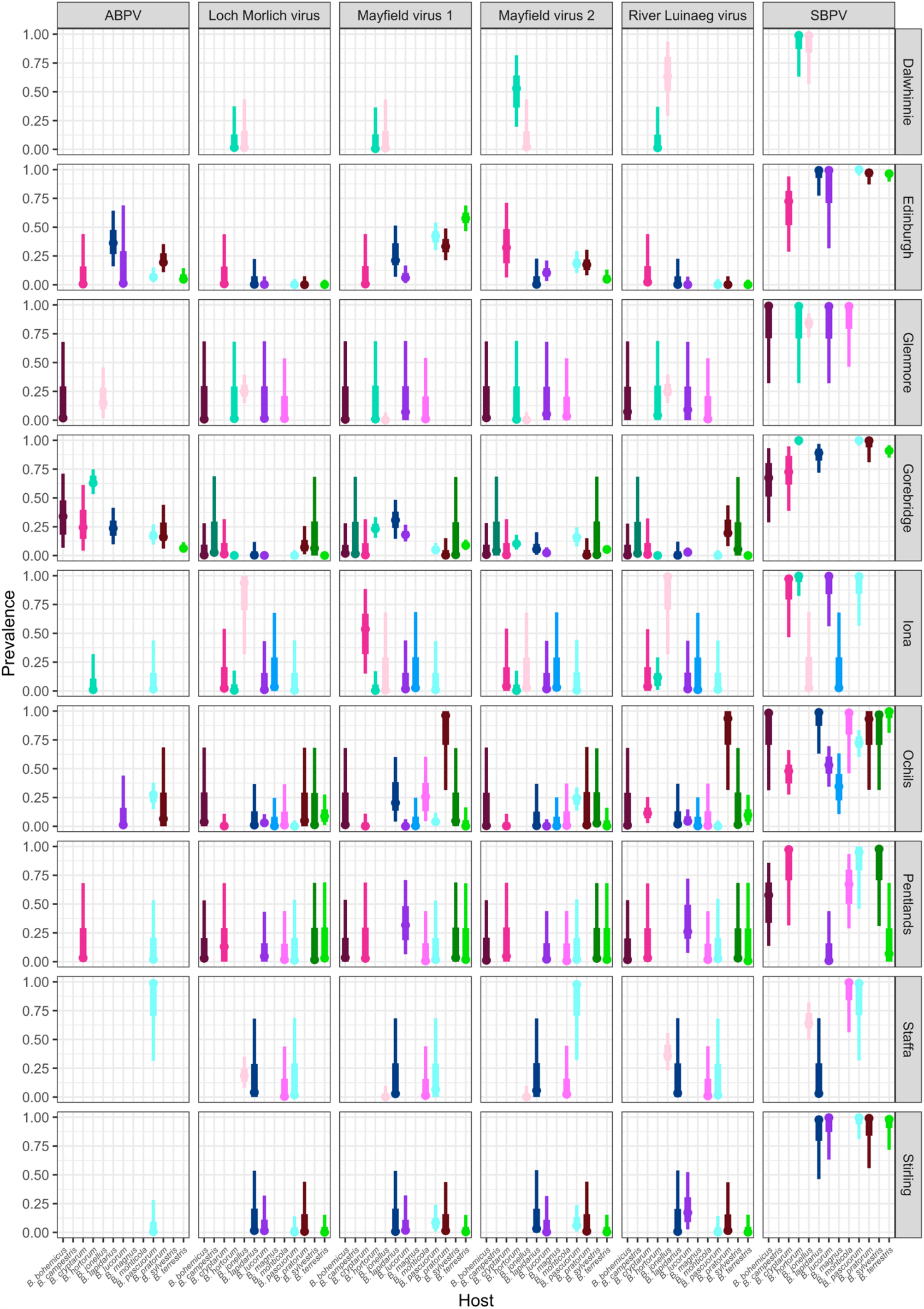
The prevalence of Acute bee paralysis virus, Loch Morlich virus, Mayfield virus 1, Mayfield virus 2, River Luinaeg virus and Slow bee paralysis virus in each sampled host species in each site. The point estimate is the posterior mode, with 50% shortest posterior intervals represented by the thick lines and 90% shortest posterior intervals represented by the thin lines. Untested combinations are left blank. Species are coloured by their corresponding colour in Figure 1 for ease of reading.

River Luinaeg virus (RLV) was detected in *B. jonellus* at all sites where the species was sampled, with prevalences of approximately 25% or higher detected at multiple sites. The prevalence was similarly high in *Bombus pratorum*. Intermediate prevalences were detected in *Bombus cryptarum*. Low levels of infection with RLV were detected in *B. lucorum* with the prevalences of the virus appearing to be considerably higher in this species in Stirling and the Pentlands. Loch Morlich virus (LMV) appears to exhibit much higher species specificity with 13/16 detections being in *B. jonellus*. It was also strongly associated with RLV, with 13/16 detections being coinfections. No species other than *B. jonellus* were detected with LMV infection in the absence of RLV coinfection. Mayfield virus 1 (MV1) appears to be a generalist, with frequent infections across bumblebee species. Its prevalence data showed large differences between the degree of infection of different bumblebee species between sites. Edinburgh and Gorebridge, two sites around 15km apart with large sample sizes, have dramatically different MV1 prevalences in *B. terrestris, B. pratorum* and *B. pascuorum*, being between 30-60% in Edinburgh, and below 15% in all species in Gorebridge. Mayfield virus 2 (MV2) shows a similar pattern but without obvious differences in infection levels between sites. The prevalence of MV2 is generally lower than that of MV1, but beyond that, the range of species infected is largely similar. Acute bee paralysis virus (ABPV) was found at intermediate modal prevalences of above 10% in all species apart from *B. terrestris* and *B. lucorum*. The prevalence of SBPV was universally high.

### Factors Influencing Infection

Viral prevalence was related to environmental covariates in some cases (Table 3; Figure 3). Higher levels of precipitation had a high posterior probability of being associated with higher prevalences of River Luinaeg virus (posterior probability: 95%), There was some evidence that more precipitation, higher maximum temperatures (which were highly correlated with solar radiation) and higher wind speeds were associated with higher prevalences of Mayfield virus 1 (posterior probability: 92%, 94% and 93% respectively). For most covariates, however, the bulk of the posterior distribution lay close to zero and did not shift considerably from the prior indicating a lack of between site resolution.

**Table 3.** The posterior correlations of the errors of each virus from the multivariate probit model, measuring the degree of co-occurance. Positive numbers represent excess co-occurrence beyond that predicted by covariates and negative numbers represent a dearth of double infections. 90% shortest posterior intervals for each correlation are shown in brackets.

|  | River Luinaeg virus | Loch Morlich virus | Mayfield virus 1 | Mayfield virus 2 | Slow bee paralysis virus | Acute bee paralysis virus |
| --- | --- | --- | --- | --- | --- | --- |
| River Luinaeg virus | 1 |  |  |  |  |  |
| Loch Morlich virus | **0.73**<br>(0.57, 0.70) | 1 |  |  |  |  |
| Mayfield virus 1 | -0.16<br>(-0.60, 0.27) | -0.17<br>(-0.69, 0.36) | 1 |  |  |  |
| Mayfield virus 2 | -0.18<br>(-0.71, 0.35) | -0.09<br>(-0.66, 0.47) | 0.05<br>(-0.13, 0.23) | 1 |  |  |
| Slow bee paralysis virus | 0.07<br>(-0.22, 0.33) | -0.06<br>(-0.36, 0.24) | 0.12<br>(-0.18, 0.41) | -0.24<br>(-0.51, 0.01) | 1 |  |
| Acute bee paralysis virus | -0.35<br>(-0.82, 0.10) | -0.31<br>(-0.81, 0.20) | 0.09<br>(-0.10, 0.28) | 0.05<br>(-0.15, 0.25) | 0.15<br>(-0.12, 0.43) | 1 |

**Figure 3.**
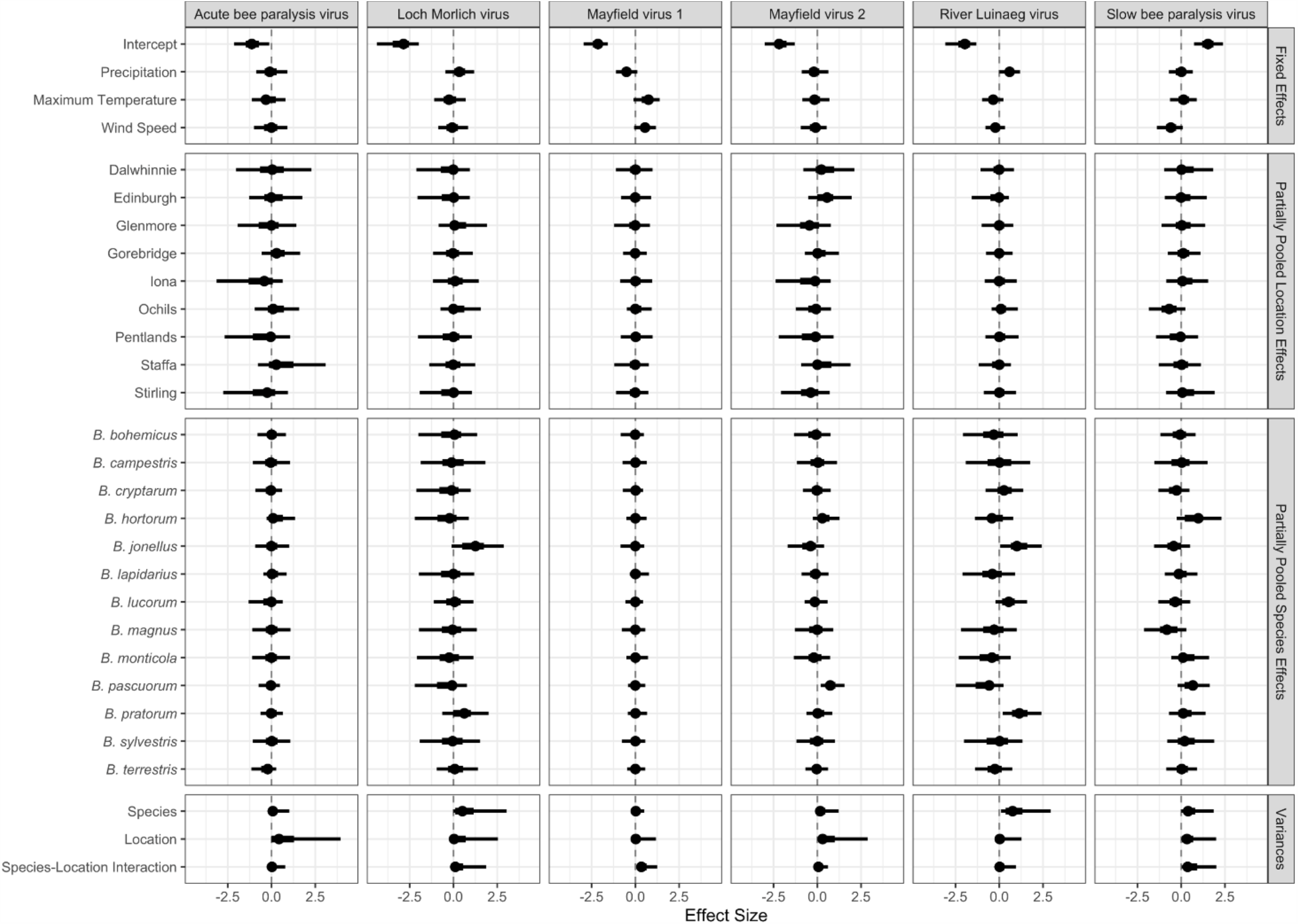
The estimates for each parameter in each virus from the multivariate probit model. The point estimate is the posterior mode, with 50% shortest posterior intervals represented by the thick lines and 90% shortest posterior intervals represented by the thin lines.

We also tested for excess co-infection beyond random between viruses using multivariate probit models that allowed us to calculate the correlation in the error terms of the multivariate normal latent variable. This measures the degree to which, after accounting for the predictors, there is still shared error, as caused by unobserved factors affecting infection risk. In this case, these measure the extent to which there is excess coinfection after accounting for the location of sampling, the species of which the bees belong and the various location-level environmental variables. Some viruses exhibited excess coinfection (Table 4). RLV andLMV showed strong and positive correlation (mean correlation: 0.73), consistent with the high levels of coinfection noted above; the error correlation between these two viruses was the only one where the bulk of the posterior was not close to zero. There was also some indication of a negative association between MV2 and SBPV, with the bulk of the posterior supporting a correlation of below zero (posterior probability: 93%), but the variance of the posterior was such that this cannot be stated with great certainty.

**Table 4.** Approximations to Watterson’s theta for sites over a homologous genomic region within the RdRp gene by host species. Combinations that weren’t tested due to low numbers of mapping reads are marked with “-“. Entries with 0.000 had no observed variation over the region under study. The point estimate is at the median predicted number of infected individuals and uncertainty corresponds to the values at the 5^th^ and 95^th^ percentile of the predicted number of infected individuals (see methods).

|  | Mixed *Bombus* pool | *Bombus terrestris*<br>pool | *Bombus lucorum* pool | *Bombus pascuorum*<br>pool |
| --- | --- | --- | --- | --- |
| River Luinaeg virus | 0.029<br>(0.029-0.030) | - | - | - |
| Loch Morlich virus | 0.026<br>(0.025-0.027) | - | - | - |
| Mayfield virus 1 | 0.010<br>(0.009-0.010) | 0.015<br>(0.014-0.015) | 0.024<br>(0.023-0.025) | - |
| Mayfield virus 2 | 0.035<br>(0.034-0.036) | 0.014<br>(0.012-0.016) | - | 0.029<br>(0.028-0.030) |
| Acute bee paralysis virus | 0.000<br>(0.000-0.000) | - | 0.000<br>(0.000-0.000) | 0.000<br>(0.000-0.000) |
| Slow bee paralysis virus | 0.008<br>(0.008-0.008) | 0.002<br>(0.002-0.002) | 0.003<br>(0.003-0.003) | 0.006<br>(0.006-0.006) |

### Diversity

Over homologous genomic regions within the RdRp gene, there were large differences between viruses in our approximation to Watterson’s theta (see methods; Table 4). RLV, LMV, MV1, and MV2 all exhibited more diversity than SBPV and ABPV, with SPBV itself being considerably more diverse than ABPV. The same genotypes of MV1 and MV2 are observed in both 2009 when Dalwhinnie, the Ochils and Iona were sampled and 2011 when Edinburgh, Gorebridge and the Pentlands were sampled, implying that the variants present in an area are stable over short periods. Additionally, as would be expected, most variation was in 3^rd^ codon positions leading to either no amino-acid replacements or replacements with similarly charged amino acids, and thus unlikely to affect protein function.

## Discussion

In this study, we explored the ecological factors influencing the genetic diversity and distribution of the viruses of wild bumblebees. We found that all the viruses detected only in bumblebees have considerably higher genetic diversity than the viruses shared with honeybees. Additionally, we found evidence of a positive association between River Luinaeg virus and Loch Morlich virus. Finally, there is some evidence for an effect of environmental variables on viral prevalence with the strongest evidence being that higher levels of precipitation increase the prevalence of River Luinaeg virus.

### Diversity

Both Acute bee paralysis virus (ABPV) and Slow bee paralysis virus (SBPV) show considerably less diversity than Mayfield virus 1 (MV1), Mayfield virus 2 (MV2), River Luinaeg virus (RLV) and Loch Morlich virus (MLV) within the study region, even though our estimates of Watterson’s theta may be conservative due to the strong assumptions on detecting viral variants in individuals. ABPV and SBPV were initially described in honeybees (Bailey et al., 1963; Bailey & Gibbs, 1964), while the other four viruses were found in bumblebees and have not been recorded in honeybees at this point (Pascall et al., 2018).Both the Sanger sequences and Watterson’s theta calculations show that the diversity in ABPV and SBPV remains low in bumblebees across the short period between 2009 to 2011. This difference could have multiple causes. It is likely that much of the observed variation in the ‘bumblebee viruses’ LMV, MV1, MV2 and RLV is neutral, as most variation is at 3^rd^ codon positions and codes for either identical or similarly charged amino acids, which are unlikely to have large fitness effects in either direction on the virus. Given the frequent bottlenecking that occurs during transmission (Zwart & Elena, 2015), such 3^rd^ codon position mutations would be inefficiently selected against.

The lack of diversity in ABPV and SBPV could be due to them infecting both honeybees and bumblebees. In multihost systems, species can differ in their susceptibility and response to infection (Ruiz-González et al., 2012), thus different species have very different levels of importance for the maintenance of a virus in a population. As such, single heavily infected host species can act as sources for infection in other sympatric species. In honeybees, these viruses interact with the mite *Varroa destructor*, a parasite that can vector viruses and is associated with the prevalence of ABPV (Mondet et al., 2014) and SBPV (Manley et al., 2020). This vector can lead to reduced viral genetic diversity in honeybees, as shown for Deformed wing virus (Martin et al., 2012), resulting in a limited pool of virus able to spill over into other hosts, which could explain the reduction in variation we observed in the viruses known to infect honeybees relative to those only described from bumblebees.

Bumblebee-limited viruses necessarily undergo multiple independent bottlenecking events when the viruses can only survive in overwintering queens, potentially maintaining diversity through reduced selection efficiency. This would initially seem to apply to SBPV, as McMahon et al. (2015) and Manley et al. (2020) report that SBPV is often found at higher prevalences in bumblebees than in sympatric honeybees during in summer. However, over winter, bumblebee populations are reduced to individual queens while honeybee colonies are maintained at population sizes in the thousands. This can maintain a level of virus in the honeybees that can then spill over into the bumblebees in the spring, which may lead to honeybee-derived SBPV variants dominating even in bumblebees. This could represent a more general effect where the fitness landscapes of viruses infecting managed species are systematically different from those infecting wild species. The hypothesised mechanism is that in a genetically homogenous, densely-packed managed population, one optimal viral genotype could easily achieve dominance as the fitness landscape will be relatively constant. On the other hand, in more genetically heterogeneous wild populations with fluctuating population sizes, the fitness landscape will be less constant, and thus selection of variants on this changing fitness landscape may lead to the maintenance of more genotypes.

### Factors Influencing Infection

We found evidence that the prevalence of River Luinaeg virus was positively associated with increased precipitation, though our ability to estimate the precise size of this effect was limited. The direction of this effect is contrary to our hypothesis that higher rainfall would decrease prevalence by reducing the contact rate between the host and virus through mechanical cleaning of contaminated floral structures and decreased bumblebee flight frequency. However, an alternative explanation consistent with the results is that high precipitation reduces foraging time and therefore condition, mediated through starvation. Starvation increases the severity of bumblebee virus infections (Manley et al., 2017) and has been shown to increase infection risk in mammals (reviewed in (França et al., 2009; Pedersen et al., 2002; Schaible & Kaufmann, 2007)). River Luinaeg virus is predominantly associated with *Bombus jonellus*, a bee species found at significantly higher frequencies in our sampling locations with higher precipitation, while the inclusion of our host random effect should account for this somewhat, it is possible that this is also driving the effect. We also found weak evidence of a negative effect of precipitation on MV1. Given that only nine sites were sampled in this study, we are limited in the between-site conclusions we can draw. Despite using a regularising prior, the risk of erroneously identifying effects can never be fully excluded in studies with small sample sizes. Thus, given the importance of wild pollinators, a study with more sites would be valuable future work to more accurately assess the effect of the environment on viral infection risk.

### Coinfection

River Luinaeg virus and Loch Morlich virus were rarely found separately in this study. They are distinct, not different segments of the same virus, as, while whole genomes are available for neither, both partial genomes include an RdRp sequence (Pascall et al., 2018). The mechanism of their transmission is unknown, but we assume that, as with most other reported bee viruses, transmission occurs at flowers.

One potential explanation for this strong association is that one of the viruses is a satellite of the other, as occurs in Chronic bee paralysis virus with Chronic bee paralysis virus satellite virus (Bailey et al., 1980). However, this seems unlikely as both virus species are observed separately, though the possibility of false negatives in the PCR reactions cannot be ruled out. Another possibility is that both viruses circulate in the population, but infection with one causes damage to the host in such a way that susceptibility to the second is dramatically increased, perhaps in a manner analogous to HIV’s synergism with TB though immune suppression (Kwan & Ernst, 2011) or influenza virus’ changing of the environment of the nasopharynx, allowing secondary bacterial invasion (Joseph et al., 2013). Viral coinfections are ubiquitously reported in prevalence studies in bees (Anderson & Gibbs, 1988; Bacandritsos et al., 2010; Blažytė-Čereškienė et al., 2016; Chen et al., 2004; Choe et al., 2012; Evans, 2001; Gajger et al., 2014; Manley et al., 2020; McMahon et al., 2015; Mouret et al., 2013; Nielsen et al., 2008; Roberts et al., 2017; Thu et al., 2016), but to our knowledge, only McMahon et al. (2015) and Manley et al. (Manley et al., 2020) tested for a departure from random expectations of infection, and no departure was found. However, non-random associations between parasites appear common, having been reported in, among other taxa including mammals (Behnke et al., 2005; Griffiths et al., 2011; Jolles et al., 2008), birds (Clark et al., 2016), arthropods (Hajek & van Nouhuys, 2016; Václav et al., 2011) and plants (Biddle et al., 2012; Seabloom et al., 2009). Thus, while the cause is uncertain, the strength of this association makes it highly unlikely to be artefactual.

## Conclusion

Here we describe the ecological and genetic characteristics of a set of viruses of bumblebees. Of the six viral species studied, genetic diversity is higher in the viruses we detected only in bumblebees. This could be explained if viruses in species managed for food production, such as honeybees, are less diverse than those in wild species, and outbreaks of these viruses in wild species are predominantly due to spillover. Further studies in this and other systems would be valuable to answer the question of whether there is a significant difference in diversity in viruses in managed species, those shared between managed and wild species, and those limited to wild species. Viral genetic diversity could be a factor in determining the risk of both disease emergence and spillover.

## Acknowledgements

We thank Dave Goulson, Claire Webster, Floh Bayer, Steph O’Connor, Penelope Whitehorn, Gillian Lye and Mario Vallejo-Marin for assistance with bumblebee collection and species identification. Additionally, we thank Angus Buckling and Andy Fenton for constructive suggestions for improvements throughout.

## Data availability statement

All code and raw infection data is available at https://github.com/dpascall/bumble_epi_diversity. All raw sequence data has been deposited on GenBank.

---

**Supporting Table 1.** The species-site breakdown of the counts of individual bumblebees used in the.

| Species | Locations |  |  |  |  |  |  |  |  | Total |
| --- | --- | --- | --- | --- | --- | --- | --- | --- | --- | --- |
|  | Dalwhinnie | Edinburgh | Glenmore | Gorebridge | Iona | Ochils | Pentlands | Staffa | Stirling |  |
| *Bombus bohemicus* | 0 | 0 | 1 | 6 | 0 | 1 | 2 | 0 | 0 | 10 |
| *Bombus campestris* | 0 | 0 | 0 | 1 | 0 | 0 | 0 | 0 | 0 | 1 |
| *Bombus cryptarum* | 0 | 3 | 0 | 5 | 2 | 19 | 1 | 0 | 0 | 30 |
| *Bombus hortorum* | 4 | 0 | 1 | 59 | 11 | 0 | 0 | 0 | 0 | 75 |
| *Bombus jonellus* | 3 | 0 | 31 | 0 | 1 | 0 | 0 | 21 | 0 | 56 |
| *Bombus lapidarius* | 0 | 8 | 0 | 17 | 0 | 4 | 0 | 1 | 2 | 32 |
| *Bombus lucorum* | 0 | 30 | 1 | 75 | 3 | 35 | 3 | 0 | 5 | 152 |
| *Bombus magnus* | 0 | 0 | 0 | 0 | 1 | 7 | 0 | 0 | 0 | 8 |
| *Bombus monticola* | 0 | 0 | 2 | 0 | 0 | 4 | 3 | 3 | 0 | 12 |
| *Bombus pascuorum* | 0 | 43 | 0 | 47 | 3 | 44 | 2 | 1 | 14 | 154 |
| *Bombus pratorum* | 0 | 29 | 0 | 13 | 0 | 1 | 0 | 0 | 3 | 46 |
| *Bombus sylvestris* | 0 | 0 | 0 | 1 | 0 | 1 | 1 | 0 | 0 | 3 |
| *Bombus terrestris* | 0 | 50 | 0 | 104 | 0 | 12 | 1 | 0 | 13 | 180 |
| Total | 7 | 163 | 36 | 328 | 21 | 128 | 13 | 26 | 37 | 759 |

**Supporting Table 2.**
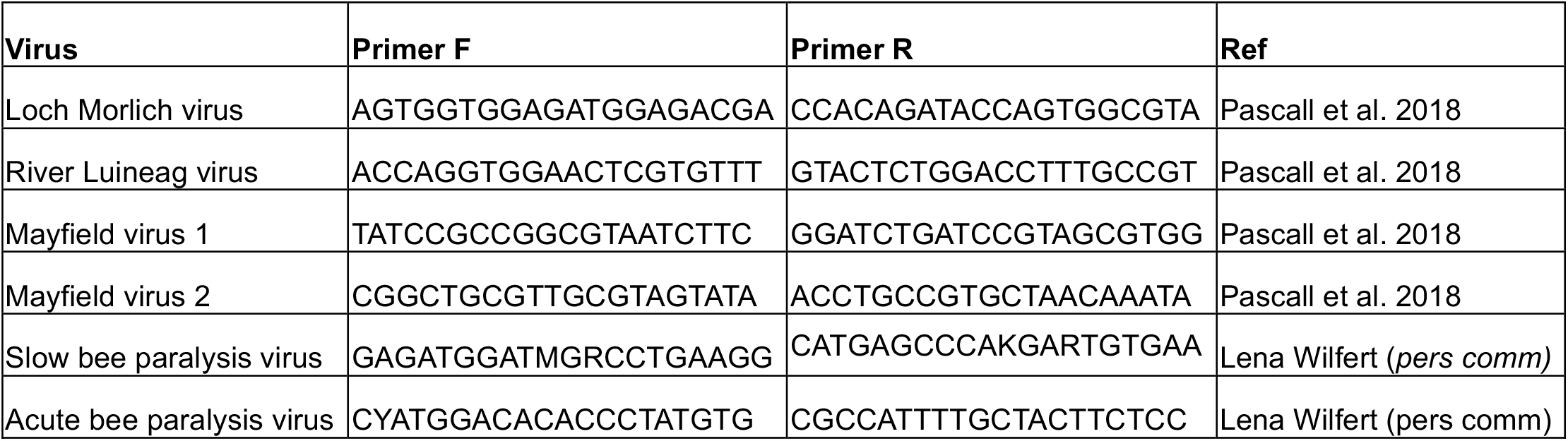
The PCR primers for each virus used in the study.

